# Characterizing protein protonation microstates using Monte Carlo sampling

**DOI:** 10.1101/2022.01.07.475457

**Authors:** Umesh Khaniya, Junjun Mao, Rongmei Wei, M. R. Gunner

## Abstract

Proteins are polyelectrolytes with acidic or basic amino acids making up ≈25% of the residues. The protonation state of all Asp, Glu, Arg, Lys, His and other protonatable residues, cofactors and ligands define each protonation microstate. As all of these residues will not be fully ionized or neutral, proteins exist in a mixture of microstates. The microstate distribution changes with pH. As the protein environment modifies the proton affinity of each site the distribution may also change in different reaction intermediates or as ligands are bound. Particular protonation microstates may be required for function, while others exist simply because there are many states with similar energy. Here, the protonation microstates generated in Monte Carlo sampling in MCCE are characterized in HEW lysozyme as a function of pH and bacterial photosynthetic reaction centers (RCs) in different reaction intermediates. The lowest energy and highest probability microstates are compared. The ΔG, ΔH and ΔS between the four protonation states of Glu35 and Asp52 in lysozyme are shown to be calculated with reasonable precision. A weighted Pearson correlation analysis identifies coupling between residue protonation states in RCs and how they change when the quinone in the Q_B_ site is reduced.

## INTRODUCTION

Proteins are large dynamic molecules that move amongst many conformations as the atoms change their positions. With N atoms there are 3N-6 vibrations ranging from high-frequency vibrations of individual bonds to larger, slower breathing modes of the protein as a whole. Positional changes can be required for function, and they are also inevitable given the low barriers for many motions. Conformational changes modify hydrogen bond patterns, expose buried charged side chains to the solution, and stabilize bound ligands.

Protons are sometimes overlooked reactants in many chemical reactions. They are lost as chemical bonds are made and many redox reactions are coupled to proton binding. The transmembrane electrochemical gradient, including a ΔpH, provides an essential store of cellular energy. Protons are therefore routinely transferred in and out of protein active sites and across membrane embedded proteins, requiring changes in protonation states along proton transfer pathways as well as at active sites. Drug binding has been shown to modify the protonation states of the protein as well as the drug itself.^1–3^ Ion binding can be coupled to changes in protonation states. ^4,5^

It has been underappreciated that proteins exist in a distribution of protonation states. An average protein has approximately 25% acidic and basic residues.^6^ A protonation microstate identifies the protonation state of all acidic and basic groups in the molecule. For N titratable groups there are 2^N^ protonation microstates. Tautomers are microstate with the same net charge but with the protons shifting position. The total number of tautomers for m protons distributed over N binding sites is

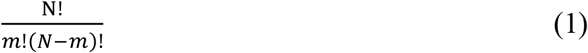

Distributions of protonation microstates occur when residue pK_a_s are near the pH in any individual structure. Also, the ensemble of protein conformations generates an additional ensemble of protonation microstates since the proton affinity of residues will shift as they sample different environments. Protonation states also change when ligands are bound^2,3^ or when redox reactions occur at bound cofactors^7,8^. Proton pumps move protons through membrane embedded proteins as changes in protein conformation and redox states modify residue proton affinity^9–13^. As with the protein conformational states, some protonation microstates are essential for protein function while others simply result from multiple microstates being close in energy.

### Analysis of microstates

The ensembles of states generated in any simulation method is limited by the degrees of freedom and the methods of moving between states. In Molecular Dynamics (MD) trajectories, atoms and molecules move in time steps, propelled by classical molecular mechanics forces. In standard MD simulations, molecules remain in a single chemical state so their protonation state is fixed at the beginning. Each time step represents a microstate of the system. In Monte Carlo (MC) sampling discrete states are generated based on the chosen degrees of freedom and then retained or rejected based on their energy. States sampled by MC can differ by changing atomic positions, number of atoms or chemical state including protonation or redox state. A microstate identifies one choice for all degrees of freedom.

MD trajectories yield large data sets with the molecule moving between many conformations. The challenge is to extract the biological significance of the data. A fraction of conformational microstates (frames), of the order of one in a thousand, are saved for analysis. Clustering algorithms group frames to cover the range of diverse conformations based on the similarity score of investigator chosen features. Well defined clustering approaches include hierarchical clustering (linkage based)^14,15^, center-based (K-means, K-centers) clustering^14,16^.

We will describe here analysis of protonation microstates obtained with the MCCE program. It uses MC sampling to generate an ensemble of residue and ligand protonation, conformation and redox states^17^. This is one of a group of programs that calculate the ensemble of protonation states for a protein using Grand Canonical Monte Carlo sampling. They find the probability of each residue being protonated at a given proton chemical potential (pH). Methods include those using semiempirical force fields^18,19^ or Continuum Electrostatics.^20,21^ They can have fixed protein positions^21,22^, include sampling side chain positions with a rigid backbone as in MCCE or as in cpHMD, be embedded in full MD.^23–27^ cpHMD usually uses a classical electrostatic force field with a vacuum dielectric constant, but can also use a polarizable force field to obtain a better treatment of the protein dielectric response^28^. Available software packages for pK_a_ prediction include MCCE^17^, PROPKA^29^, DelPhiPka^30^, H^++31^, PypKa.^32^

Monte Carlo sampling requires consideration of millions of randomly chosen microstates. Traditional calculation of protonation using MC compresses the output to provide only the average probability for the protonation state of each residue at a given pH. A titration is simulated by carrying out the calculation at multiple pHs to derive a pK_a_s for all residues. If other degrees of freedom, such as ligand binding or residue conformation are allowed, their probabilities are also found. However, the full microstate distribution contains additional information such as the range of protein net charge and the correlation between protonation of individual sites and with any other available degrees of freedom. Microstate analysis can find the probability of higher energy states that may be reaction intermediates. The protonation microstates can also provide a complete assignment of protonation states for all residues and tautomer states for all His as input for MD. The microstates with the lowest energy or highest probability can be compared.

Here we will describe using Grand Canonical Monte Carlo (GGMC) within MCCE to characterize the distribution of protonation microstates and their thermodynamic properties using hen white lysozyme and *Rb. sphaeroides* photosynthetic reaction centers (RCs) as examples. Lysozyme has been used as a test case for pK_a_ predications and the protonation probability as a function of pH has been measured.^33–36^ RCs have a complex network of protonatable residues that function to modulate the electrochemistry of quinone reduction and to serve as a proton transfer pathway to the quinone.^37,38^ This larger system will show the complexity of protonation microstates in a system that requires proton transfer for function. The microstate energy distribution, the distribution of microstates with unique charges, the thermodynamics of an individual protonation reaction and the correlation of the protonation of individual residues will be described.

## METHODS

The Boltzmann distribution of protonation states and side chain polar proton positions are found for hen egg-white Lysozyme (PDB ID: 4lZT^39^) and the reaction centers (RCs) from the purple non-sulfur photosynthetic bacteria *Rb. Sphaeroides* (PDB ID: 1AIG^40^) using the MCCE program^17^.

### Conformer generation determines the degrees of freedom

In MCCE the protein backbone is always fixed. Residue side chains and ligands can be given multiple choices that define their protonation and conformational state. Each choice is a ‘conformer’. While MCCE can carry out full side chain rotamer sampling, ‘isosteric’ conformers are used here for simplicity^41^. MCCE can also include explicit waters with multiple positions.^42^ However as is customary, all water molecules in the crystal structure are deleted and replaced by implicit solvent in lysozyme while 28 crystallographic waters are retaining in the RCs. All protonatable residues (Glu, Asp, Arg, Lys, His, Tyr and Cys and N- and C-termini) have conformers that are charged and others that are neutral. Side chain conformers are made that differ in the position of polar side chain protons. This includes conformers with different neutral His tautomers, Asn and Gln amide termini orientation and the position of hydroxyl protons on Tyr, Ser, Thr and Cys and proton positions on neutral Asp and Glu. The ubiquinone in RCs can be oxidized, Q, or anionic semiquinone, Q^•-^. Each residue has from one to six conformers that are selected in MC sampling.

### MCCE force field

A microstate of the protein is one selected conformer for each residue. The energy of each microstate is divided into a reference energy, and self and pairwise terms.^17^ The energy of any microstate is obtained from the pre-calculated energy lookup table. The energy (ΔH^x^) of microstate x is:

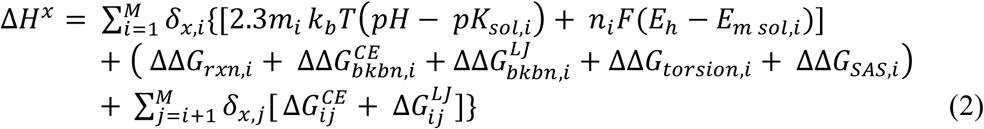

The first line describes the energy for each residue type to be in its protonated or unprotonated form or to be oxidized or reduced in solution. Lines two and three will describe how the protein shifts the proton and electron affinity of the conformer. M is the total number of conformers. *δ*_x,i_ is 1 if conformer i is present in microstate x or 0 otherwise. m_i_ is 1 for basic protonated, -1 for acidic deprotonated and 0 for all neutral conformers. pK_sol,i_ is the reference pK_a_ of this residue type in solution, E_m_ is the electrochemical midpoint potential of this redox active group. F is the Faraday constant, n_i_ is the change in the number of electrons on a conformer type that is taking part in a redox titration. pH and E_h_ are the relevant solution parameters describing the chemical potential of protons or electrons that will come to equilibrium with the protein.

The second line of equation 2 describes the self-energies of the conformer that form the microstates. These energies are independent of the conformer choice for other residues. These are: the loss of conformer solvation energy as it is moved from solution to its position in the protein, torsion energy, continuum electrostatic (CE) and Lennard Jones (LJ) van der Waals interactions with the fixed backbone amides and favorable, van der Waals interactions of the exposed side chain surface with the implicit solvent. The third line gives the continuum electrostatic and Lennard Jones pairwise interactions between each pair of conformers in other residues. All energies are calculated prior to MC sampling. Electrostatic energies are calculated using the Poisson-Boltzmann solver in Delphi^20,43,44^ using Parse charges^45^, a dielectric constant of 4 for protein and 80 for solvent with an implicit salt concentration of 150 mM. MCCE corrects the continuum electrostatic energies for the changes in the dielectric boundary due to changes in surface rotamers.^17^ Non-electrostatic energies use the Amber force field.^46^

### Monte Carlos sampling

**In MCCE** microstates change by random choices, first of a residue then of a conformer available for that residue. If the chosen residue contains conformers which interact with an absolute value >0.5 kcal/mol with conformers of another residue, additional conformer changes for that other residue will be made 50% of the time. Here conformers in several residues switch, followed by a single decision using the Metropolis–Hastings algorithm to accept or reject the change. Allowing one to three residues to change helps avoid electrostatic or van der Waals clashes between strongly coupled groups and speeds convergence.^47,48^

The Monte Carlo sampling routine starts with a random microstate with one conformer assigned to each residue. The program then goes through annealing, conformer reduction, and MC sampling. Annealing changes the acceptances temperature in the Metropolis-Hastings acceptance criteria. MC sampling is then carried out at room temperature. By default, equilibration samples 3000 MC steps/conformer for lysozyme and 300 for RCs. Prior to the production phase, conformers that are rarely chosen are removed from the list of sampled conformers. The default cut-off used here, is that a conformer with a probability less than 0.001 is discarded. This speeds convergence and results in acceptance of MC steps of ≈30%. However, the early elimination of conformers creates problems in highly correlated systems so it can be modified as needed.^42^ The discarded conformers are excluded from all later sampling. “Free residues” have a choice of conformers. A residue whose protonation state becomes fixed by all conformers of one protonation type being discarded may still sample conformers with different positions. If the probability of a single conformer for a residue is unity then that is a fixed residue, which is removed from the sampling list.

The energy for the full ensemble is calculated once for the first random microstate. In each subsequent step the interactions with the conformer chosen for change is subtracted from the ensemble energy and the energy of the new conformer for that residue is added back, speeding the energy calculation in each MC step. The default number of Monte Carlo steps for the production phase, which microstates will be recorded, is 2000 * number of free conformers. Six independent Monte Carlo cycles of this length starting with a random microstate formed from free conformers are carried out. The six runs are combined, but they can be compared to check the quality of Monte Carlo convergence.

### Microstate information storage

A major challenge is to record the several million accepted microstates in a readable form. The MCCE algorithm has several features that make this easier. One is that residue conformers are premade and discrete so that we can use the conformer indices as a unique identifier for each microstate. Also, only a few residues are changed on each step so that we need only record the conformers that have changed when a microstate is accepted. Also, no dynamic information needs to be recorded for fixed residues as their conformer never changes.

The microstate file starts with making a list of conformer IDs that are removed from sampling because they had low probability during pre-production MC sampling and the list of conformers that will be sampled. Likewise, a list of the fixed residues with only one available conformer is separated from the list of free residues which have multiple available conformers. The conformers for the fixed residues are saved at the start. The first line of each MC run gives the initial microstate, listing the occupied conformer ID for all free residues. Subsequent lines record only the conformer IDs that have changed in this step, the microstate energy (kcal/mol) and the microstate count, which is the number of microstates that are rejected before the next, new microstate is accepted. Links to Jupiter notebooks with tools and tutorials for microstate analysis are found in SI Link S1.

### Generating the protonation microstate ensemble

The microstate file is a ticker-tape that must be replayed to find the subsequent microstates. The information can be sorted on many properties. Here we will describe how protonation microstates are identified. Many conformational microstates are aggregated for each unique protonation microstate.

First, the charge of fixed residues that have only a single conformer available in MC is determined. This charge is summed and treated as the background charge which will be added to find the total protein charge in each microstate.

Then the ticker-tape is read to place all microstates of free residues into memory. The microstate file gives the conformer ID of each free residue. A microstate is constructed with only the charge of this conformer. This is a vector of the length of all free residues with an entry of -1, 0 or 1 (though groups with other charge states are allowed). Then we determine which charge microstates are unique. The length of the unique microstate vectors is reduced to contain only residues of interest, here acidic and basic residues and the ubiquinone. At each stage the MC count is retained and is summed when multiple microstates are grouped into one category.

### Correlation between microstates

Correlation measures the strength of the relationship between two variables with the sign giving the direction of the trend. The weighted Pearson Correlation (r_pq_)^49^, used here to find residues whose protonation changes are coupled together, is given by:

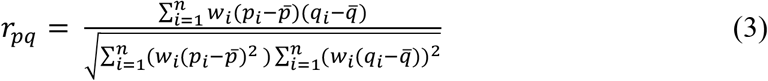

Here *w*_*i*_ is the weight, p_i_ and q_i_ are the protonation states for two residues in a microstate i, 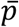 and 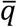 are the mean value of p_i_ and q_i_ respectively and n is the number of unique accepted charge microstates. The unique charge file is the input. Only those residues whose protonation states take different values in the ensemble are included. The weight of each microstate is the count of accepted microstates that have this unique protonation state. r_pq_ is calculated for all pairs of residues with variable charge.

## RESEULTS AND DISCUSSION

Traditionally MCCE analysis provides the Boltzmann averaged protonation and side chain and ligand position for a protein^41^ or other macromolecules.^50,51^ The goal is to now characterize the distribution of protonation microstates in the Boltzmann distribution. In MCCE a microstate defines both residue and ligand charge and position. Protonation microstates, which define the charge of every acidic and basic residue, will exist in many conformational states. The charge state identifies the net, total charge in the microstate. Tautomers are groups of protonation microstates that have the same charge but with the protons distributed over different residues.

### Microstate Energy distribution for lysozyme

Lysozyme is a small protein with 129 residues that is a benchmark protein for pK_a_ calculations.^17^ The seven Asp, two Glu, one His, six Lys and N- and C-termini are assigned ionized and neutral conformers. The isosteric conformer routine without explicit waters in MCCE created 284 conformers. Approximately 1 to 1.5 million microstates are sampled. Considering both protonation and conformation degrees of freedom 94% of the accepted microstates in Monte Carlo sampling are unique, meaning they are not revisiting a previously accepted microstate.

Figure 1 shows the distribution of microstate energies in the Boltzmann distribution. Even this small protein, there is a significant energy range. The full width at half maximum (FWHM) of the skewed normal distribution is 3.61 kcal/mol. There are a small number of microstates at lowest energy, which are well separated from those at highest probability. The shape of the probability distribution is similar at all pHs and for small proteins such as lysozyme and large proteins such as RCs (Figure S1).

**Figure 1.**
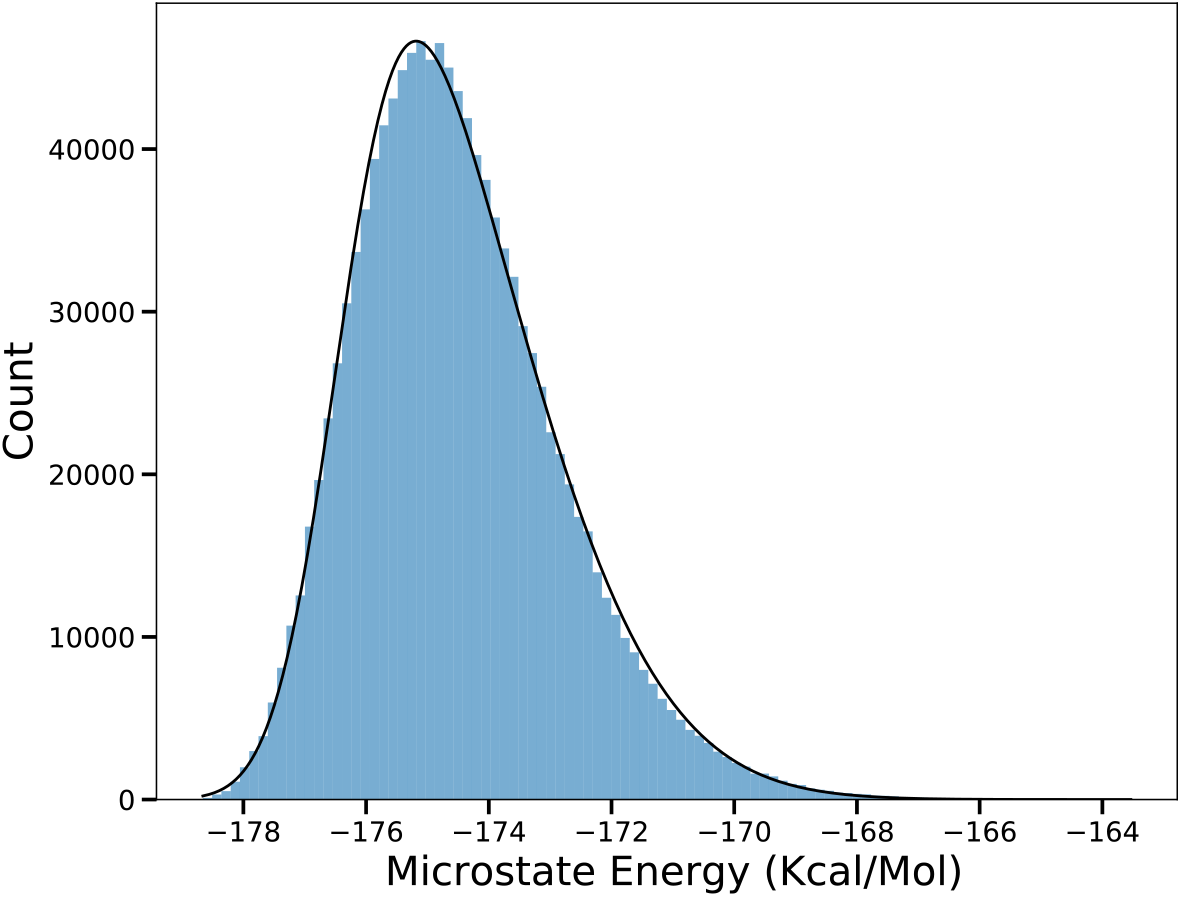
Microstate Energy distribution for lysozyme (PDB ID: 4LZT) at pH 7. The count is the number of times microstates in that energy window are accepted. Black line is best fit to a skewed normal distribution curve with skew of 2.86.

In the lysozyme calculation, at pH 4 there are >500,000 unique accepted conformation/protonation microstates and >200,000 at pH 7. Near the residue pK_a_ both protonated and deprotonated conformers are in the accepted ensemble. Thus, the number of unique accepted microstates reflects the number of residues that are titrating (Table 1). In lysozyme most residues have pK_a_s close to their solution values. Thus, at low pH the acids are titrating. At pH 4 ten residues have significant probability of being either charged or neutral leading to 222 unique protonation microstates. Near pH 7 most Asp and Glu and the C-terminus are stably deprotonated and Lys and Arg protonated. Here, the protonation microstates reflect the mixture of His and N-terminus protonation states (Table S1). At pH 7 only six residues are in a distribution of protonation states and there are only 24 different protonation microstates in the accepted ensemble (Table 1).

**Table 1.** Characterization of accepted microstates obtained with Metropolis-Hastings sampling. The MC steps are the aggregate of 2,000 times the number of free conformers times six restarts. Unique microstates have different protonation and or conformation. Protonation microstates differ in location of protons on acidic and basic groups. *Number of unique protonation states within 1.36 kcal/mol of the lowest or highest energy or of ±0.68 kcal/mol of the average energy. There are 27 protonatable residues (Asp, Glu, Arg, His and Lys) and chain termini in lysozyme and 132 in RCs. The number of residues that change protonation have different charge states in the microstate ensemble. MC sampling for RCs is carried out with the ubiquinone in the QB site being the neutral quinone, QB, the anionic semiquinone QB•^-^ or with the Eh at the Em for the quinone so there is a 50:50 mixture of the two states.

|  | Number: All microstates |  | Number: Protonation microstates |  |  |  | Number: Residues |
| --- | --- | --- | --- | --- | --- | --- | --- |
|  | MC steps | Unique accepted (%) | Unique | Lowest energy* | Average energy* | Highest energy* | Change protonation |
| pH | Lysozyme |  |  |  |  |  |  |
| 4 | 1,500,000 | 527,014 (95.4%) | 222 | 4 | 89 | 6 | 10 |
| 5 | 1,500,000 | 484,152 (94.59%) | 147 | 2 | 28 | 2 | 11 |
| 6 | 1,200,000 | 430,904 (94.28%) | 38 | 2 | 15 | 7 | 7 |
| 7 | 1,200,000 | 447,522 (94.72 %) | 24 | 4 | 16 | 4 | 6 |
| State | RCs (pH 7) |  |  |  |  |  |  |
| Q<sub>B</sub> | 8,700,000 | 2,735,799 (99.19%) | 60, 381 | 3 | 24,254 | 1 | 53 |
| 50:50 | 8,294,680 | 2,765,120 (99.19 %) | 70,713 | 4 | 27,775 | 6 | 51 |
| $Q_{B^{\bullet-}}$ | 8,700,000 | 2,720,450<br>(99.21%) | 54,448 | 5 | 22,202 | 3 | 48 |

### The distribution of charge and tautomer states

Figure 2 shows the distribution of the unique accepted protonation microstates. This mirrors the distribution of charge states that can be found experimentally. However, experimental measurements of protein net charge combine the charge of the protein and of bound ions^52–54^. There are 147 unique protonation states for lysozyme at pH 5 and 24 at pH 7. The ensemble average charge is 9.68 at pH 5 and 7.78 at pH 7 (Figure S2). At pH 5 proteins with charge ranging from 7 to 13 are found, though those with charge 9 or 10 are most probable, having the highest count. Each vertical line of dots is a group of tautomeric states, with the same charge but different proton distributions. In MCCE every unique charge microstate exists in a many tautomer and conformational microstates covering a wide energy range (Figure S3). Larger and more yellow dots in Figure 2 have more underlying conformational states with this distribution of protons.

**Figure 2.**
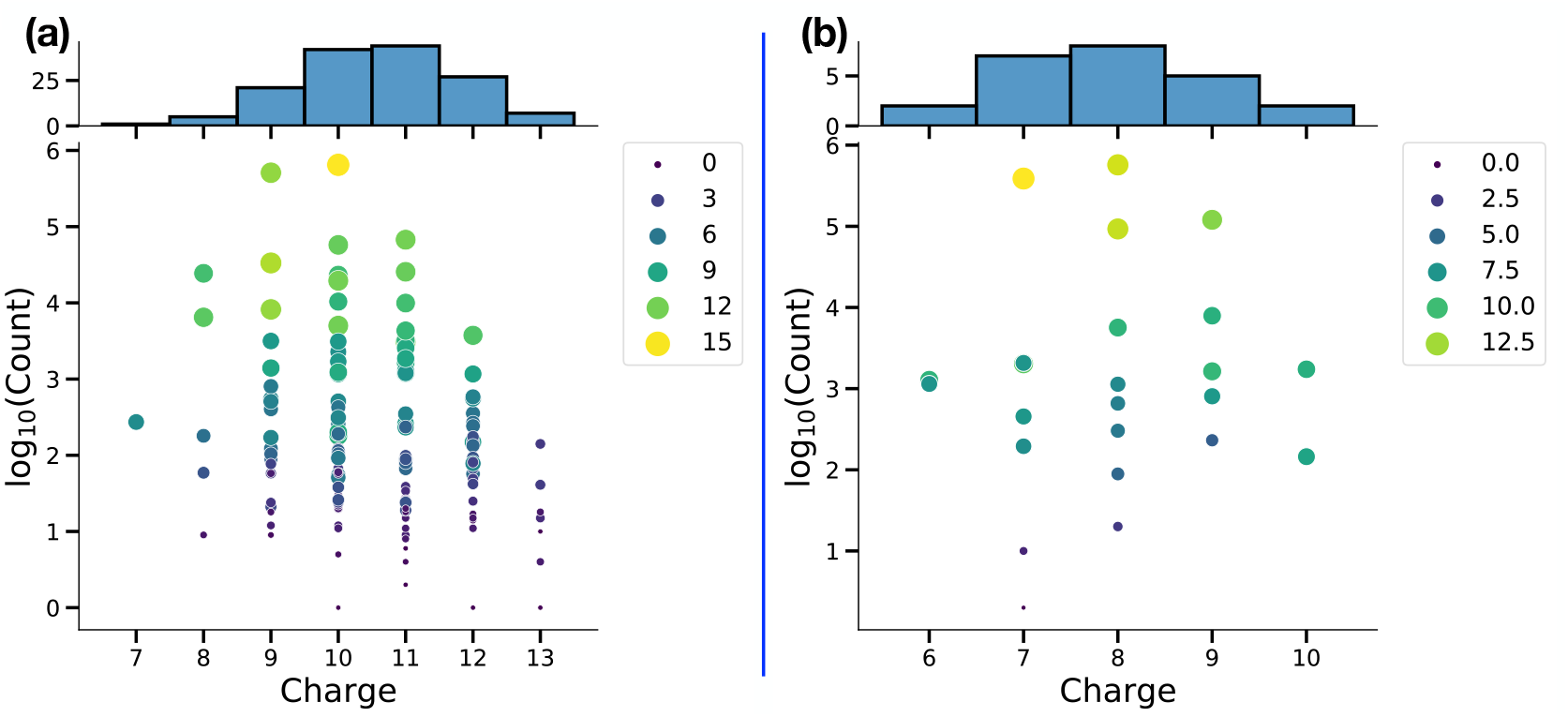
Unique tautomer charge distribution for lysozyme at pH (A) 5 and (B) 7. Each point in the scatter plot is a unique protonation microstate. The count gives its acceptance in MC sampling, with high count indicating higher probability. Color and size of points correspond to the range of microstate energies associated with all microstates found for that protonation microstate. Histogram at the top shows the total number of unique protonation microstates with a given charge. Thus, it counts the number of tautomers at each charge.

Even a system as small as lysozyme can have a significant number of unique protonation microstates. However, only a small number are found with significant probability in the Boltzmann ensemble. Thus, there are 2^18^ possible protonation states, but at pH 7 only 6 residues have different protonation states in the microstate ensemble. However, of the 2^6^ (64) possible states only 24 are ever in an accepted microstate. Figure 2 shows there are two microstates at either pH 5 or 7 that have several fold more occupancy in MC sampling than any others. Table S1 lists all 24 accepted protonation microstates for lysozyme at pH 7 as an example. Only three microstates have a probability greater than 10% and they sum to 90% of the ensemble. At pH 5, 7 states sum to a probability of 90%.

### Thermodynamic parameters for protonation reactions from analysis of the microstate ensemble

The changes in the free energy of a reaction determines the proportion of reactants and products at the system at equilibrium (K_eq_). The changes in enthalpy and entropy give insight into the nature of what changes in the reaction. Measurements of the equilibrium constant, K_eq_, ΔH and ΔS are common, and there are methods to calculate these parameters from MD trajectories.^55–57^ However, calculating ΔG, ΔH and especially ΔS for complex biological molecules often leads to large uncertainty as it can be hard to sample enough states.^58–62^ Here we analyze the accepted microstates in MCCE to determine the thermodynamic properties of the coupled protonation reactions of Glu35 and Asp52 in lysozyme (Figure 3).

**Figure 3.**
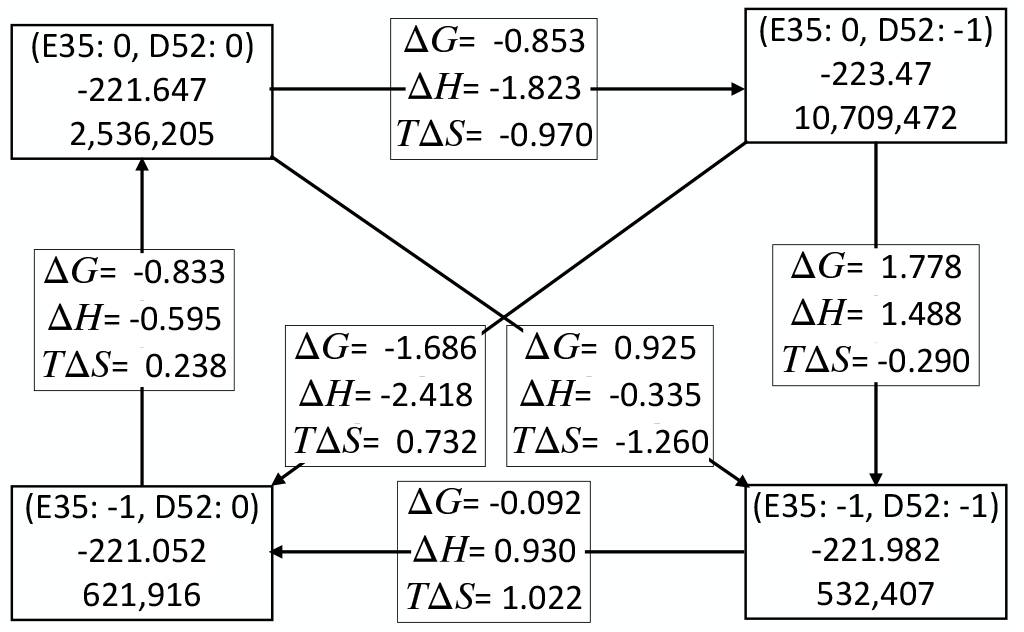
Thermodynamic box for titration of Glu35 and Asp52 in lysozyme at pH 4 at 298.15 K. The full microstate ensemble is sorted into the four protonation states of these two residues and these groups are independently characterized. Corner boxes: first row is charge of Glu35 and Asp52; second row is the average microstate energy (H); third row is the count of microstates in this protonation state. The boxes along the arrows give the ΔG, ΔH and TΔS for the transition between different protonation states. The top right and bottom left states are tautomers. All energies are in kcal/mol.

**Figure 4.**
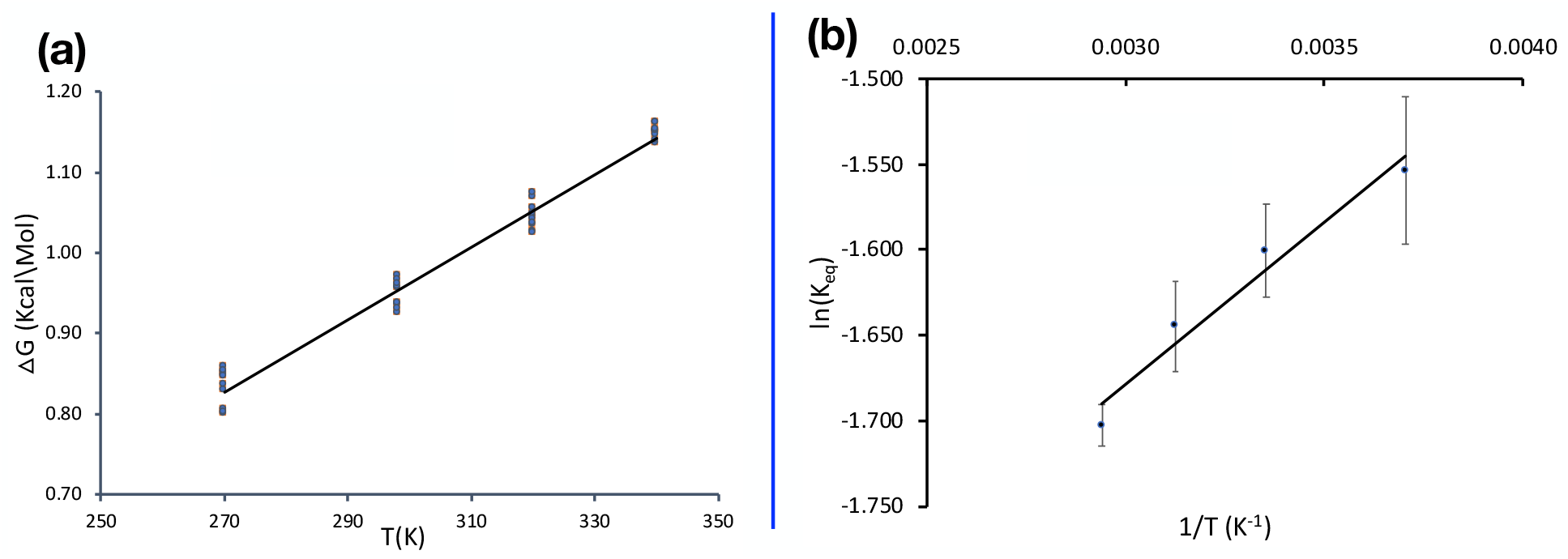
Temperature dependence of thermodynamic parameters. (A) Change in Δ*G*for the reaction taking Glu35 and Asp52 with both neutral to both ionized with temperature. Line is the linear best fit and each dot represent the Δ*G*value for an independent MC sampling run using the same input conformer energies. The slope of the graph gives the -ΔS of 0.0045 kcal/mol/° and intercept gives the ΔH of 0.387 kcal/mol, R^2^ is 0.979. (B) Van’t Hoff plot for the same dataset. The slope is ΔH/R and intercept ΔS/R where R is the gas constant. ΔH is 0.377 kcal/mol and -ΔS is 0.0045 kcal/mol.

To determine the thermodynamic variables, the MC microstates are divided into four groups, each with a different protonation state for the two residues. The enthalpy is the average energy for the ensemble of microstates with appropriate protonation for the two acids:

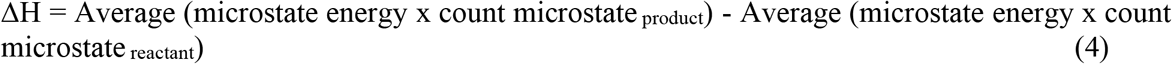

The change in free energy between pairs of states is:

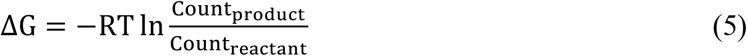

Therefore, entropy is:

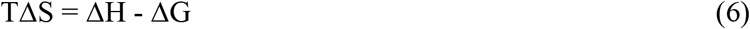

Additional MC sampling reduces the standard deviation for the calculation of ΔS. Here the number of MC steps for annealing and for the production phase are increased 10-fold, with 20,000 steps/conformer carried out with the sampling of the free conformers restarted with a new randomly chosen microstate six times.

Figure 3 shows the results for one MCCE calculation at pH 4 where the system is in a mixture of the four protonation microstates. The results for the reaction starting from both acids neutral to both ionized was calculated for 10 independent MC calculations gives values of ΔG of 0.95±0.16 kcal/mol with a ΔH of -0.35±0.02 and -TΔS of 1.30±0.02 kcal/mol. Thus, within the constraints of the limited degrees of freedom of the MCCE calculation thermodynamic variables can be derived from the Boltzmann ensemble with reasonable precision.

Examination of the thermodynamic square shows that the state with Glu35 neutral and Asp52 ionized is favored. This agrees with the ensemble average probability where Glu35 is 8% ionized and Asp52 is 78% ionized. The breakdown of ΔH and ΔS shows that ΔH plays the larger role, favoring this state.

### Thermodynamic properties with change in temperature

The ΔS is traditionally measured from the temperature dependence of a reaction. The same can be done here. The same energy look-up tables are used, but the temperature of 270 K, 298.15 K, 320 K and 340 K is used for the Metropolis-Hastings test for microstate acceptance. Ten independent MC calculations are carried out at each temperature. All calculated ΔG, ΔH and ΔS for the system are provided in the Table S3B-D. The standard deviation of the thermodynamic variables is <0.2 kcal/mol, showing the MCCE microstate ensemble can provide reproducible values even for the estimate of ΔS. The same values are obtained from the Van’t Hoff plot of K_eq_ vs 1/T.^59^ The reaction is exothermic so heat would be released on ionization of the two acids. The standard deviation of the ΔG or K_eq_ is larger at lower temperatures.

### Correlation analysis of protonation states of residues in RCs

A major advantage of having access to the microstates it that the correlation between conformer choices can be found. Here we will describe the correlation of residue protonation states in different microstates using the much larger RCs as an example. It has 828 residues in the coordinate file. There are 132 ionizable residues (Asp, Glu, Arg, Lys and His) and NTR, CTR, and two redox active quinones and other cofactors. There are 1743 isosteric conformers for the protein, cofactors and retained water molecules. More than 45 residues are found in a mixture of ionization states at pH 7 and there are more than 50,000 unique protonation microstates to investigate (Table 1).

RCs show the distinction between the lowest energy microstates and those with highest probability. We determined the lowest energy example of each of the thirty most probable protonation microstates (Figure S4). These were compared with the ranking of the states at lowest energy. While the high probability states are enriched with lower energy microstates they are not identical. This shows that the conformational degrees of freedom can create more opportunities to make some protonation microstates than others.

In this system the proton distribution is functionally important as a proton must be brought into the Q_B_ site, 15Å from the surface, through a network of protonatable residues and waters.^63–^ _65_ The Pearson weighted correlation coefficient is used to find the correlation between the residues (equation 3). Residues whose charge state is the same in all microstates are removed from the analysis, as these are not correlated with any other. The weighted correlation coefficient is obtained by taking the unique charge microstates with its corresponding count for the ≈45 remaining residues. Only the residues that have an absolute value of the correlation of at least 0.1 with any other residue are included in the heatmap (Figure 5). This procedure identifies 15-16 residues, all of which are near the Q_B_ site.

**Figure 5.**
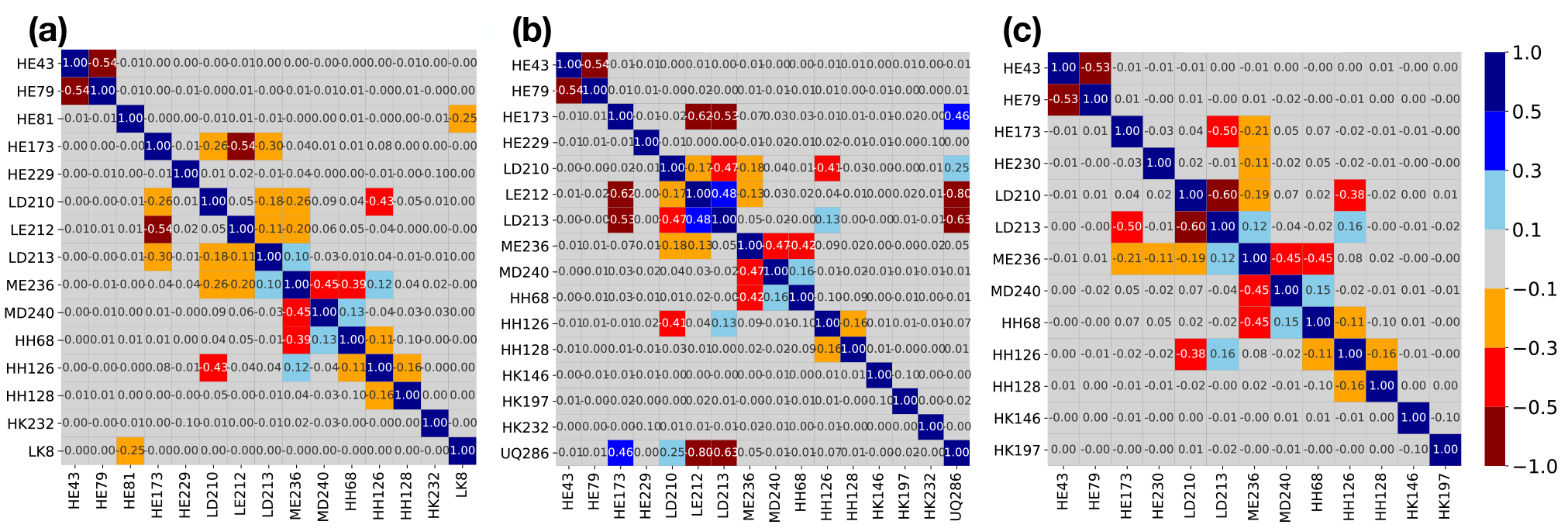
Heat map of Pearson’s weighted correlation coefficient of residue ionization states in RCs at pH 7. Only residues with an absolute value for the correlation ≥0.1 with at least one other residue are shown. Each square in the heat map gives the correlation strength for the two residues obtained with equation 3. Residue are identified as: chain (L, M or H), one letter residue name, then residue number. Ubiquinone is UQ. Dark blue, blue, sky blue, light gray, orange, red and dark red correspond to the correlation value of range 0.5 to 1.0, 0.3 to 0.5, 0.1 to 0.3, -0.1 to 0.1, -0.3 to -0.1, -0.5 to -0.3 and -1.0 to -0.5 respectively. (a) Neutral quinone (b) 50:50 mixture of two state (c) Anionic semiquinone QB•^-^. The quinone is not seen in panel a or c as it is 100% in a single redox state.

The correlation analysis carried out for the RC protonation microstate distribution in three quinone states: neutral quinone, ≈50:50 mixture of the two redox states and the anionic semiquinone Q_B_•^-^. A positive correlation indicates that the two groups are more likely to be ionized together, while negative correlation shows that ionization of one reduces ionization of the other. Thus, in the presence of neutral Q_B_ (Fig. A) the ionization of the group of acidic residues, GluH79, AspL210, GluL212 and AspL213 are negatively correlated with each other. Ionization of AspM240 and HisH68 are positively correlated. In the presence of Q_B_•^-^ the heat map has two fewer residues than it does in the presence of Q_B_. GluL212, which is very close to the quinone, is now always protonated.^7^ As the quinone redox state is expected to modify the protonation states in its vicinity, a heat map is prepared at an E_h_ where the quinone is ≈50% reduced in the MCCE MC sampling. The protonation of residues that are correlated with the quinone redox state can be seen. Thus, GluL212 and GluL213 are less ionized when the quinone is reduced, while GluH173 and AspL210 are more ionized. Figure 6 shows the position of the residues that are interacting with each other.

**Figure 6.**
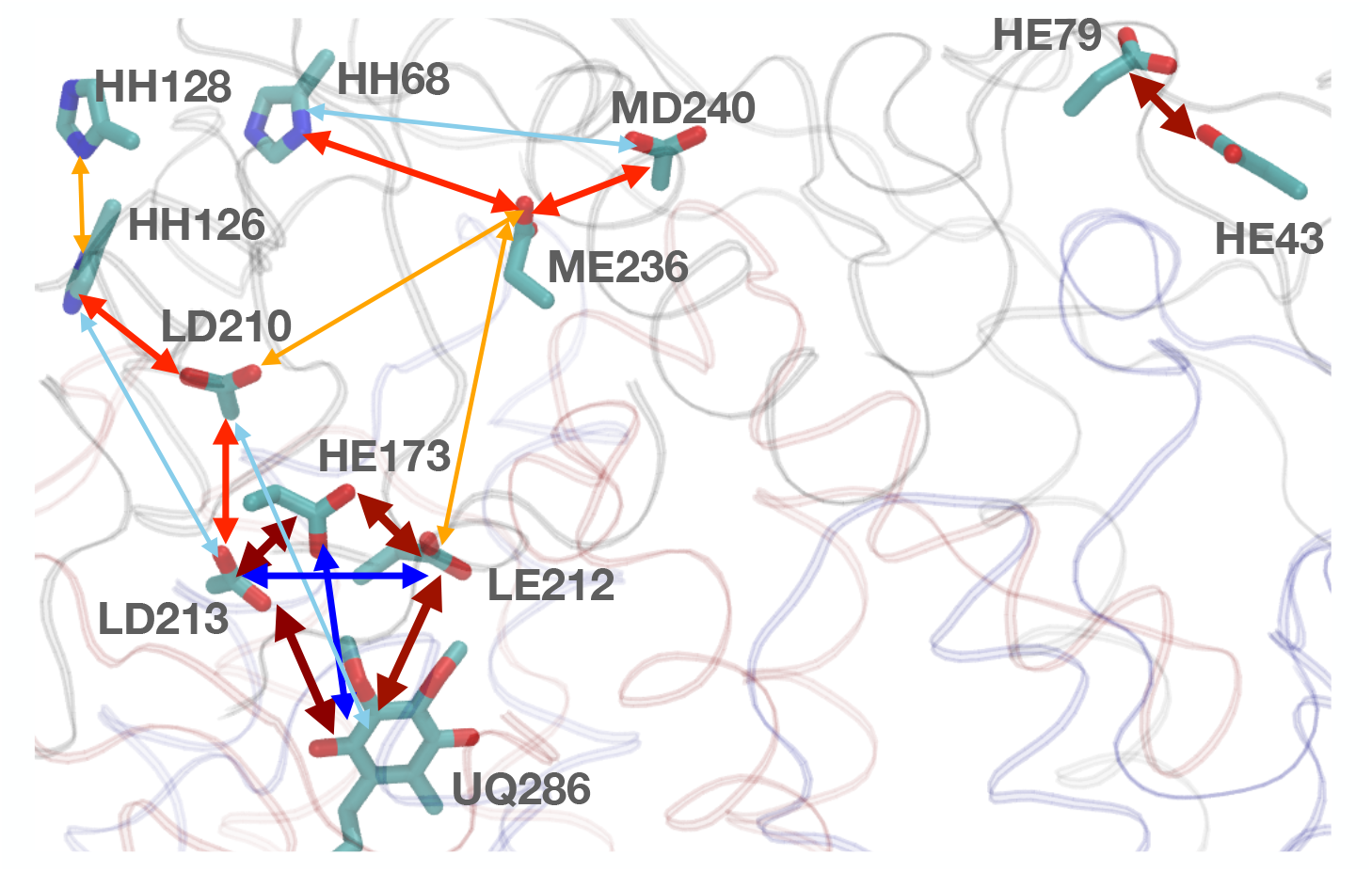
Structure representation of residues that show correlations in the MC sampling with a 50:50 mixture of QB and Q_B_•^-^. Only those residues whose absolute correlation coefficient is ≥0.1 are shown. Blue and red two-sided arrows show the positive and negative correlation respectively. The color code is same as Figure 5. Wider arrows indicate higher correlation.

### Other uses of microstate analysis in MCCE

There are other important properties of proteins whose analysis relies on knowing the microstates rather than simply the average properties. The correlation of structures and protonation state can show how small changes influence the proton affinity of interior sites in the protein.^66^ In proton pumps these changes can lead to efficient proton loading and unloading as protons are transferred through the protein.^12,67^ Microstate analysis can show how the protonation states of ligands can shift those of the protein. The proton transfer paths require continuous hydrogen bonded connections.^68,69^ MCCE has been used to identify the hydrogen bonds in the MC ensemble to trace proton transfer paths.^42,70–72^ High probability or low energy microstates that define residue protonation and neutral His tautomers provide rational input to standard MD.

## CONCLUSIONS

Proteins have many acidic and basic residues and are very unlikely to be in a single protonation state. Microstate analysis allows us to characterize the ensemble of protonation microstates. This shows proteins have a range of charge states and for each charge state that there are multiple tautomers with different distribution of protons. While there can be an astonishing number of protonation microstates found in proteins such as RCs only a few have significant probability. The lower population states may play roles as intermediates in important processes such as proton transfers. The microstate analysis shows how the protonation state of groups of residues are coupled together and which residues are not correlated with each other. Analysis, not carried out here, can show how individual hydrogen bonding patterns can stabilize particular protonation states or be aligned for proton transport. The MC ensemble is large enough that calculations of the thermodynamic parameters including the ΔS of proton binding can be calculated with reasonable precision.

## ACKNOWLEDGMENTS

The authors would like to acknowledge the funds from the National Science Foundation grant MCB-1519640.

## Table of Contents Image

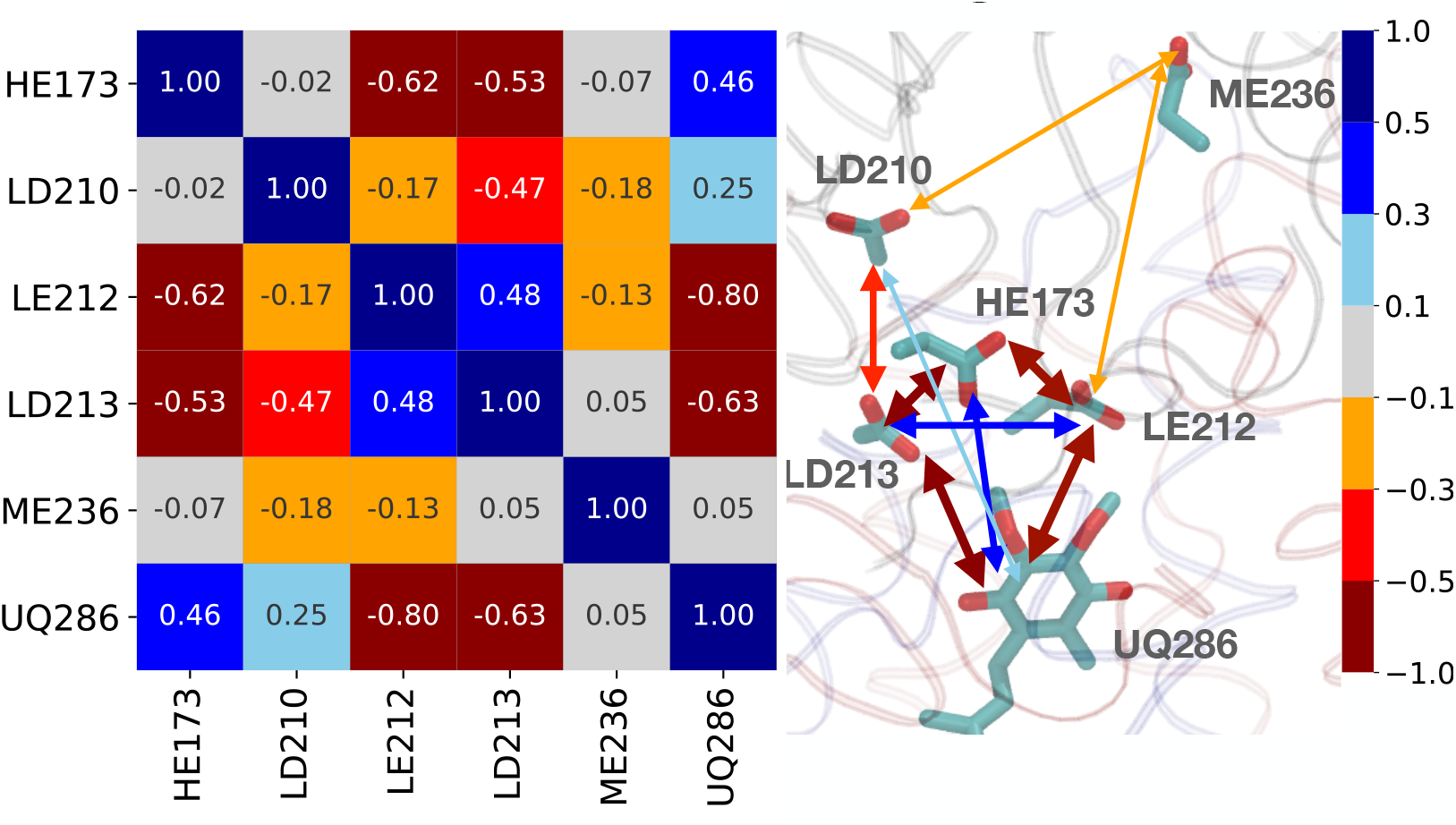

Figure: Table of content image

