## Supplementary file for "Characterizing protein protonation microstates using Monte Carlo sampling"

**Figure S1**. Microstate Energy distribution for different pH, different temperature and different proteins.

**Figure S2**. Average ensemble protein charge of Lysozyme from pH 4 to pH 7.

**Figure S3**. Microstate Energy distribution for each charge of 4lzt at pH 5 and pH 7

**Figure S4.** Correlation between the probability of a protonation microstate and its energy ranking in RCs at pH 7 with neutral Q_B_.

**Table S1**. Occupancy and charge state of individual unique charge microstate of residues that are changing the ionization state in Monte Carlo simulation.

**Table S2**. Reproducibility of values characterizing the microstate ensembles.

**Table S3.** Reproducibility of thermodynamic parameters.

**Link S1.**: Links to the programs used here.

**Figure S1**. Microstate Energy distribution for different pH, different temperature and different proteins.

Black line is skew normal distribution fitted curve of histogram.

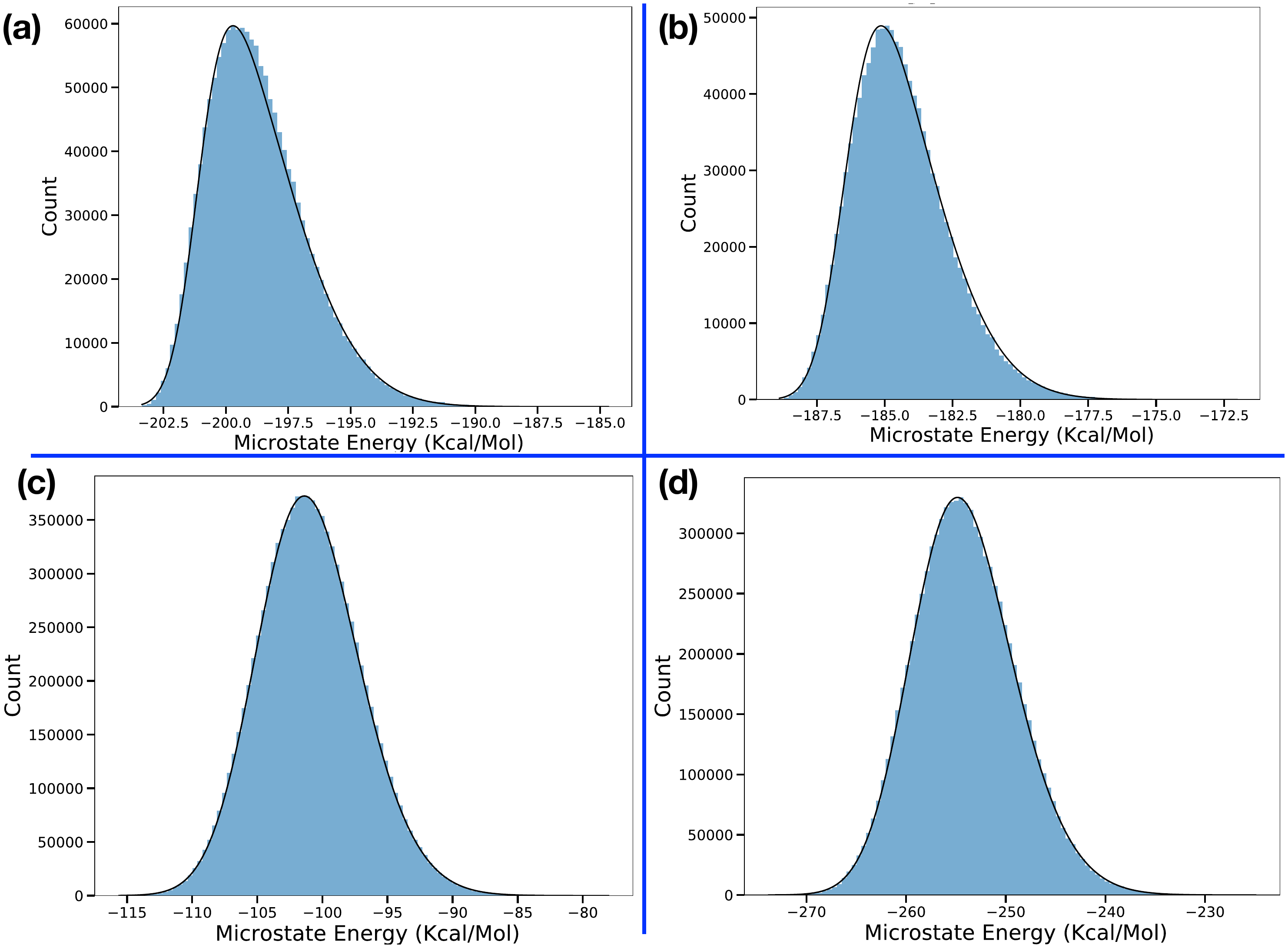

(a) Lysozyme at pH 5 at T = 298.15 K, skew 3.41, FWHM 4.22 kcal/mol.

(b) Lysozyme at pH 7 at T = 320 K, skew 2.90, FWHM 3.86 kcal/mol.

(c) Lysozyme at pH 7 at T = 298.15 K (default run), skew 1.28, FWHM 9.19 kcal/mol.

(d) RCs at pH = 7 at T = 298.15 with neutral quinone, skew 1.47, FWHM 11.89 kcal/mol.

**Figure S2**. Average ensemble protein charge of Lysozyme from pH 4 to pH 7.

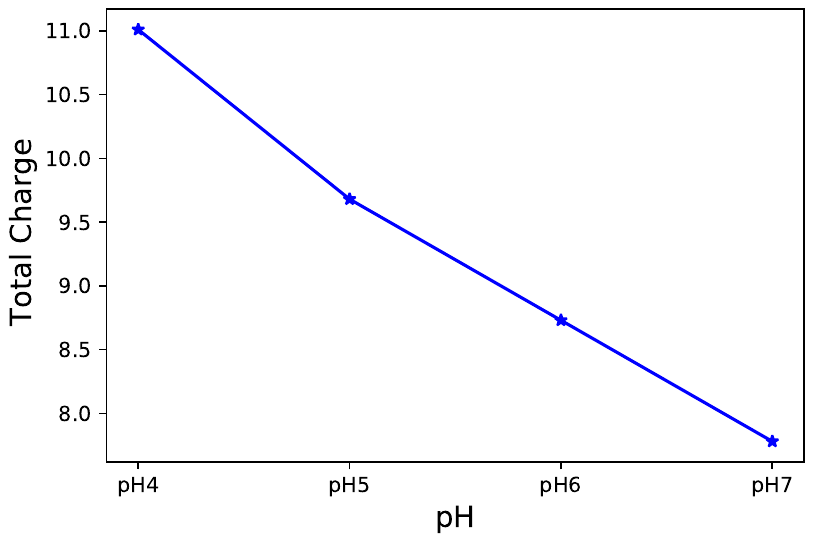

The average charge changes from 11.01 at pH 4 to 7.78 at pH 7. At each pH the protein has a non-integer average charge indicating that there are acidic and basic residues in a mixture of protonation states in the ensemble.

**Figure S3**. Microstate Energy distribution for each charge of 4lzt. (a) pH 5 (b) and pH 7

The wide energy range for each protonation microstate arises from the large number of conformational microstates available to most protonation microstates. Thus, even at pH 7 where there are only 8 tuatomers with a charge of 8, there are 249,209 total microstates with that charge.

**Figure S4**. Correlation between the probability of a protonation microstate and its energy ranking in RCs at pH 7 with neutral Q_B_.

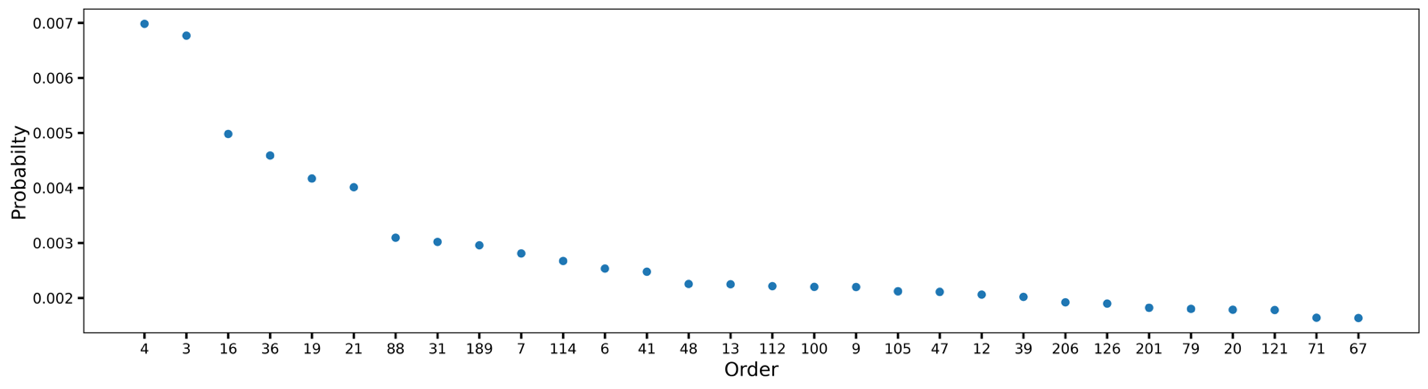

The probability of the 30 microstates with the highest probability. The x axis notes the order of energy of the conformational microstate with this distribution of protons that has the lowest energy. Thus, the protonation state of the microstate with the lowest energy found in MC sampling (i.e. order 1) is not one of the highest probability protonation microstates. While there are over 50,000 unique protonation microstates, these 30 protonation microstates account for 8.48% of the total population of the ensemble**.**

**Table S1**. Occupancy and charge state of individual unique charge microstate of residues that are changing the ionization state in Monte Carlo simulation.

Only three microstates have occupancy ≥10%. They account for 90% of the total population.

| **NTR1** | **K1** | **H15** | **E35** | **D101** | **K116** | **Count** | **Net Charge** | **Occupancy in Fraction** |
| --- | --- | --- | --- | --- | --- | --- | --- | --- |
| 0 | 1 | 1 | -1 | -1 | 1 | 571852 | 8 | 0.477 |
| 0 | 1 | 0 | -1 | -1 | 1 | 387654 | 7 | 0.323 |
| 1 | 1 | 1 | -1 | -1 | 1 | 120227 | 9 | 0.100 |
| 1 | 1 | 0 | -1 | -1 | 1 | 92767 | 8 | 0.077 |
| 0 | 1 | 1 | 0 | -1 | 1 | 7910 | 9 | 0.007 |
| 0 | 1 | 0 | 0 | -1 | 1 | 5654 | 8 | 0.005 |
| 0 | 1 | 1 | -1 | -1 | 0 | 2073 | 7 | 0.002 |
| 0 | 0 | 1 | -1 | -1 | 1 | 2024 | 7 | 0.002 |
| 1 | 1 | 1 | 0 | -1 | 1 | 1732 | 10 | 0.001 |
| 1 | 1 | 0 | 0 | -1 | 1 | 1634 | 9 | 0.001 |
| 0 | 1 | 0 | -1 | -1 | 0 | 1286 | 6 | 0.001 |
| 0 | 0 | 0 | -1 | -1 | 1 | 1142 | 6 | 0.001 |
| 1 | 0 | 1 | -1 | -1 | 1 | 1133 | 8 | 0.001 |
| 0 | 1 | 1 | -1 | 0 | 1 | 805 | 9 | 0.001 |
| 0 | 1 | 0 | -1 | 0 | 1 | 658 | 8 | 0.001 |
| 1 | 0 | 0 | -1 | -1 | 1 | 454 | 7 | 0 |
| 1 | 1 | 1 | -1 | -1 | 0 | 303 | 8 | 0 |
| 1 | 1 | 0 | -1 | 0 | 1 | 231 | 9 | 0 |
| 1 | 1 | 0 | -1 | -1 | 0 | 195 | 7 | 0 |
| 1 | 1 | 1 | -1 | 0 | 1 | 145 | 10 | 0 |
| 0 | 1 | 1 | 0 | -1 | 0 | 89 | 8 | 0 |
| 0 | 0 | 1 | 0 | -1 | 1 | 20 | 8 | 0 |
| 0 | 1 | 0 | 0 | -1 | 0 | 10 | 7 | 0 |
| 1 | 0 | 1 | -1 | -1 | 0 | 2 | 7 | 0 |

**Table S2**. Reproducibility of values characterizing the microstate ensembles.

| pH | Structure | Summary | Unique  Charge MS | Number of Unique Charge MS | | | # residues or termini changing protonation |
| --- | --- | --- | --- | --- | --- | --- | --- |
|  |  |  |  | Lowest energy* | Average energy* | Highest energy* |  |
| pH4 | 4lzt | Av | 285.50 | 4.30 | 98.80 | 5.80 | 11.10 |
|  |  | STD | ±36.27 | ±0.67 | ±5.98 | ±4.64 | ±0.74 |
| pH5 | 4lzt | Av | 117.30 | 2.00 | 30.00 | 5.30 | 10.22 |
|  |  | STD | ±13.43 | ±0.00 | ±2.71 | ±3.43 | ±0.67 |
| pH6 | 4lzt | Av | 43.90 | 2.10 | 15.60 | 4.80 | 7.50 |
|  |  | STD | ±6.03 | ±0.32 | ±1.17 | ±2.53 | ±0.71 |
| pH7 | 4lzt | Av | 28.90 | 4.00 | 15.00 | 4.30 | 6.80 |
|  |  | STD | ±4.48 | ±0.00 | ±1.05 | ±2.58 | ±0.79 |
| RCs | | | | | | | |
| pH7 | Q_B_ | Av | 59,734.20 | 6.60 | 24,021.20 | 2.60 | 51.60 |
|  |  | STD | ±1,891.18 | ±2.45 | ±662.45 | ±1.82 | ±3.13 |
| pH7 | 50:50 | Av | 70,139.40 | 8.00 | 27,411.60 | 5.00 | 50.80 |
|  |  | STD | ±1,614.04 | ±5.48 | ±357.08 | ±4.18 | ±1.92 |
| pH7 | Q_B_•^-^ | Av | 5,4795.80 | 3.00 | 22,203.80 | 2.60 | 49.40 |
|  |  | STD | ±932.91 | ±1.41 | ±472.35 | ±1.67 | ±3.44 |

Characterization of accepted microstates obtained with Metropolis-Hastings sampling. The MC steps are the aggregate of 2,000 times the number of free conformers times six restarts. Unique microstates have different protonation and or conformation. Protonation microstates differ in location of protons on acidic and basic groups. *Number of unique protonation states within 1.36 kcal/mol of the lowest or highest energy or of ±0.68 kcal/mol of the average energy. There are 27 protonatable residues (Asp, Glu, Arg, His and Lys) and chain termini in lysozyme and 132 in RCs. The number of residues that change protonation have different charge states in the microstate ensemble. MC sampling for RCs is carried out with the ubiquinone in the Q_B­_ site being the neutral quinone, Q_B_, the anionic semiquinone Q_B_•^-^ or with the E_h_ at the E_m_ for the quinone so there is a 50:50 mixture of the two states.

**Table S3.** Reproducibility of thermodynamic parameters.

The reaction is for the two residues E35 and D52 of lysozyme at pH 5 with both neutral as the reactant and both ionized as the product. The ∆G, ∆H and ∆S are calculated from the MC ensemble using equations 4, 5 and 6 in main text. $\Delta G$ units are in kcal/mol and temperature in Kelvin unit.

| Table S3A | Average energy (kcal/mol) | | | |
| --- | --- | --- | --- | --- |
| Temp. (K) | **270.00** | **298.15** | **320.00** | **340.00** |
| run_1 | -223.722 | -222.99 | -222.374 | -221.794 |
| run_2 | -223.731 | -222.996 | -222.363 | -221.791 |
| run_3 | -223.740 | -222.977 | -222.364 | -221.808 |
| run_4 | -223.717 | -222.988 | -222.381 | -221.785 |
| run_5 | -223.728 | -222.963 | -222.359 | -221.794 |
| run_6 | -223.726 | -222.996 | -222.353 | -221.816 |
| run_7 | -223.749 | -222.970 | -222.348 | -221.794 |
| run_8 | -223.747 | -222.969 | -222.375 | -221.808 |
| run_9 | -223.740 | -222.981 | -222.361 | -221.814 |
| run_10 | -223.723 | -222.972 | -222.368 | -221.777 |
| **AV** | **-223.732** | **-222.980** | **-222.365** | **-221.798** |
| STD | ±0.011 | ±0.012 | ±0.010 | ±0.013 |

| Table S3B | Average ∆G (kcal/mol) | | | |
| --- | --- | --- | --- | --- |
| Temp. (K) | **270** | **298.15** | **320** | **340** |
| run_1 | 0.852 | 0.925 | 1.045 | 1.136 |
| run_2 | 0.829 | 0.936 | 1.042 | 1.139 |
| run_3 | 0.800 | 0.957 | 1.026 | 1.152 |
| run_4 | 0.837 | 0.931 | 1.025 | 1.152 |
| run_5 | 0.859 | 0.938 | 1.035 | 1.15 |
| run_6 | 0.849 | 0.938 | 1.070 | 1.161 |
| run_7 | 0.806 | 0.958 | 1.074 | 1.147 |
| run_8 | 0.802 | 0.971 | 1.048 | 1.152 |
| run_9 | 0.847 | 0.966 | 1.056 | 1.153 |
| run_10 | 0.854 | 0.962 | 1.037 | 1.161 |
| **Av ∆G** | **0.834** | **0.948** | **1.046** | **1.150** |
| STD | ±0.023 | ±0.016 | ±0.017 | ±0.008 |

| Table S3C | ∆H (kcal/mol | | | |
| --- | --- | --- | --- | --- |
| Temp. (K) | **270** | **298.15** | **320** | **340** |
| run_1 | -0.242 | -0.335 | -0.418 | -0.469 |
| run_2 | -0.266 | -0.353 | -0.432 | -0.482 |
| run_3 | -0.240 | -0.313 | -0.383 | -0.476 |
| run_4 | -0.247 | -0.356 | -0.409 | -0.531 |
| run_5 | -0.257 | -0.367 | -0.486 | -0.495 |
| run_6 | -0.266 | -0.348 | -0.459 | -0.488 |
| run_7 | -0.276 | -0.376 | -0.444 | -0.522 |
| run_8 | -0.294 | -0.326 | -0.397 | -0.482 |
| run_9 | -0.279 | -0.366 | -0.438 | -0.498 |
| run_10 | -0.241 | -0.354 | -0.417 | -0.546 |
| **Av ∆H** | **-0.261** | **-0.349** | **-0.428** | **-0.499** |
| STD | ±0.019 | ±0.020 | ±0.030 | ±0.026 |

| Table S3D | **T∆S (kcal/mol)** | | | |
| --- | --- | --- | --- | --- |
| Temp. (K) | **270** | **298.15** | **320** | **340** |
| run_1 | -1.094 | -1.260 | -1.463 | -1.605 |
| run_2 | -1.095 | -1.289 | -1.474 | -1.621 |
| run_3 | -1.040 | -1.270 | -1.409 | -1.628 |
| run_4 | -1.084 | -1.287 | -1.434 | -1.683 |
| run_5 | -1.116 | -1.305 | -1.521 | -1.645 |
| run_6 | -1.115 | -1.286 | -1.529 | -1.649 |
| run_7 | -1.082 | -1.334 | -1.518 | -1.669 |
| run_8 | -1.096 | -1.297 | -1.445 | -1.634 |
| run_9 | -1.126 | -1.332 | -1.494 | -1.651 |
| run_10 | -1.095 | -1.316 | -1.454 | -1.707 |
| **Av T∆S** | **-1.094** | **-1.298** | **-1.474** | **-1.649** |
| STD | ±0.024 | ±0.024 | ±0.040 | ±0.030 |
| Av ∆S  Cal/mol/° | 0.004 | 0.004 | 0.005 | 0.005 |

**Link S1.** Links to the programs used here.

Programs used here can be found on GitHub.

1. The MCCE program is available at: <https://github.com/GunnerLab/Stable-MCCE>
2. Microstate analysis Library tutorial: https://gunnerlab.github.io/Stable-MCCE/ms_analysis/
3. Jupyter Notebook tutorial for microstate analysis: https://github.com/umeshkhaniya/ms_tutorial
